# PLATE: a global atlas of food metaplasmidomes

**DOI:** 10.64898/2026.11.03.763617

**Authors:** Mehmet Umur Aslan, Rıza Arda Kayacan, Süleyman Arif Söğütdelen, Özlem Sağıroğlu, Muzaffer Arıkan

## Abstract

Plasmids play a central role in the acquisition and spread of genes underlying important traits in food-associated microorganisms. Although shotgun metagenomics has greatly expanded opportunities to study food microbiomes, their plasmid diversity remains largely uncharacterized. Here, we present the Plasmid Atlas of Food Ecosystems (PLATE), a curated global resource of food metaplasmidomes. Built from 4,690 food metagenomes spanning 55 countries and 17 categories, PLATE comprises a broad catalog (bPLATE) of 3,904,895 putative plasmid contigs (PPCs) and a refined catalog (rPLATE) of 213,474 PPCs, corresponding to 181,871 unique sequences. Both catalogs are complemented by curated sample metadata, host predictions, taxonomic assignments, and functional annotations covering antimicrobial resistance (AMR), protein families and KEGG orthologs, CRISPR-Cas systems, carbohydrate-active enzymes, and bioactive peptides. We demonstrate PLATE’s utility through a case study of AMR genes detected on PPCs from food microbiomes. PLATE is freely accessible and enables users to interactively search for and download sequences, annotations, and associated metadata. By organizing plasmid-associated sequence diversity and functional potential across food ecosystems, PLATE provides a foundation for analyses of food metaplasmidomes.

## INTRODUCTION

Microorganisms associated with foods influence fermentation, product characteristics, spoilage and safety (1–4). Plasmids in food-associated bacteria can carry genes relevant to fermentation and microbial adaptation, including those involved in lactose and citrate utilization, proteolysis, exopolysaccharide and bacteriocin production, stress tolerance, and antimicrobial resistance (5–8). These traits can influence growth, competitiveness and technological performance in food environments. Therefore, understanding the genetic content of food-associated plasmids is important for characterizing the functional potential of food microbiomes.

Most knowledge of plasmid-encoded traits comes from cultured isolates, leaving their distribution across food microbial communities poorly understood. Food metagenomics has expanded access to microbial diversity across food types and geographic regions, including microorganisms that are difficult to cultivate (9). By recovering putative plasmid contigs (PPCs) directly from community DNA, metagenomic approaches also enable investigation of the metaplasmidome without requiring cultivation of individual hosts. Thus, large-scale studies have applied these approaches to characterize plasmid diversity and predicted functions in soil, aquatic and other environments (10, 11). Extending these efforts to food ecosystems requires linking PPCs to their sample origins, food categories and functional annotations so that their genetic content can be examined alongside their distribution across foods.

Several public databases provide complementary resources for plasmid research. PLSDB (12) curates complete plasmid sequences from nucleotide archives, providing reference sequences for plasmid identification and comparison. IMG/PR (13) expands coverage through systematic prediction of plasmids from isolate genomes, metagenomes and metatranscriptomes within IMG/M, accompanied by functional annotations and ecological metadata. PlasmidScope (14) integrates sequences from multiple repositories and provides standardized annotations, interactive visualization and online analytical tools. These resources support broad investigations of plasmid diversity, but their coverage across environments does not provide a dedicated atlas of plasmids recovered from food metagenomes. A resource that systematically organizes food-derived plasmid candidates using consistent sequence-selection criteria, harmonized food categories and traceable sample provenance is needed to examine plasmid-associated genetic potential across food ecosystems.

Here, we present the Plasmid Atlas of Food Ecosystems (PLATE), a dedicated resource integrating PPCs recovered from 4,690 food-associated metagenomes with harmonized sample metadata and consistent functional annotations. PLATE comprises a broad catalog (bPLATE) and a more stringently filtered refined catalog (rPLATE). We characterize rPLATE’s sequence composition, representation in existing plasmid resources, predicted hosts and functional potential across food categories, and demonstrate its utility through a sequence-based case study of recurrent AMR gene combinations.

PLATE is freely accessible at https://www.plate-db.com, where users can explore PPCs, inspect linked annotations and sample provenance, and download data for further analysis. By connecting putative plasmid content with food-source information, PLATE supports comparative investigations across food ecosystems and the selection of candidates for experimental studies.

## MATERIALS AND METHODS

### Data acquisition, sample processing and food classification

Food-associated shotgun metagenomic datasets were obtained from NCBI (n = 645), the curated Food Metagenomic Database (cFMD; n = 3,990) (1) and MGnify (n = 53) (15). NCBI records were identified using the following query: [“food” AND “shotgun” AND “metagenome” AND “biomol DNA” AND “library layout paired” AND “platform Illumina”]. Records were restricted to genomic libraries, and sample annotations were reviewed to exclude non-food metagenomes, including wastewater and industrial datasets. Food metagenomes were retrieved from cFMD without additional source-specific filtering. MGnify datasets were selected from the food production biome, excluding amplicon datasets and samples unrelated to food. Overlapping records across sources were deduplicated to obtain 4,688 public datasets.

Two additional datasets, kefir and boza, were generated in-house. DNA was extracted using the Stool DNA Isolation Kit (Norgen Biotek) according to the manufacturer’s instructions and sequenced on an Illumina NovaSeq platform with 150-bp paired-end reads. The combined dataset comprised 4,690 datasets.

Preassembled contigs for 4,586 public datasets were obtained from Logan (16). The remaining 102 public datasets and two in-house datasets were assembled from raw reads following the same pipeline applied for preassembled contigs. Briefly, reads were quality-checked using FastQC (https://www.bioinformatics.babraham.ac.uk/projects/fastqc) and MultiQC (17), and filtered using Trimmomatic (18). Unitigs were generated using Cuttlefish2 (19) and subsequently assembled into contigs using Minia3 (20).

Datasets were assigned to 17 source categories using a modified cFMD classification scheme: alcohol, dairy, fermented beverages, fermented grains, fermented fruits and vegetables, fermented legumes, fermented meat, fermented seeds, fermented tubers and roots, seafood, fruits and vegetables, meat, probiotics, feed, supplements, water and other. The feed, supplements and water categories were added to the original scheme, and fish, fermented fish and other seafood-related categories were combined into seafood.

### Plasmid prediction and construction of the catalogs

Putative plasmid contigs (PPCs) were identified with Plasmer (21) using default parameters, excluding sequences shorter than 500 bp or longer than 500,000 bp. The complete set of Plasmer-positive sequences constituted the broad catalog (bPLATE), comprising 3,904,895 PPCs.

A more stringent refined catalog (rPLATE) was constructed through sequential filtering. bPLATE PPCs were first restricted to sequences ε2 kb and screened using PlasClass (22). PPCs with a plasmid probability >0.5 were retained and evaluated using viralVerify (23). Sequences classified as Virus or Chromosome were removed. The remaining sequences were classified using PlasFlow (24), chromosome and unclassified calls were excluded, and only plasmid calls were retained. Ribosomal RNA-positive sequences were identified using Rfam (25) models and Infernal (26), following the previously described workflow (10), and excluded. This subset was then screened against the human reference genome GRCh38 using minimap2 (27). PPCs with alignments exceeding 95% nucleotide identity and 80% query-contig coverage were removed. Sequence redundancy within the resulting rPLATE catalog was assessed by identifying identical sequences using SeqKit (28) and determining the number of sequence clusters using MMseqs2 (29) with thresholds of 95% nucleotide identity and 85% coverage of the target (cluster-member) sequence. bPLATE and rPLATE were retained as separate resources, with catalog membership being distinguished in downstream summaries.

To assess the reliability of plasmid prediction, PLSDB plasmids and cFMD prokaryotic MAGs were fragmented into random 2–50 kb segments with matched length distributions between the two classes. Prior to fragmentation, MAG contigs aligning to PLSDB plasmids (minimap2 -x asm10; ≥50% query coverage and ≥90% identity) were removed. Fragment lengths were drawn from a single uniform distribution shared by both classes, fragments were taken from a random strand, and plasmids were treated as circular sequences. Three replicates of 5,000 fragments per class were generated with independent random seeds. All fragments were classified with Plasmer using default parameters, and the default class call (P(plasmid) ≥ 0.5) was taken as the bPLATE prediction. The rPLATE refinement steps were then applied to bPLATE plasmid calls as in the main pipeline. The human-genome filter was omitted because the benchmark contained no human sequences. For comparison, PlasClass and PlasFlow were also applied independently to all fragments. Recall, specificity and false positive rate (FPR) were calculated from the confusion matrices.

### Comparison with existing plasmid databases

Unique rPLATE PPCs were compared against PLSDB (12), PlasmidScope (14) and IMG/PR (13) using minimap2 (27). Nucleotide identity was calculated as the number of matching bases divided by the alignment block length, and query coverage as the aligned query length divided by the full contig length. Coverage was evaluated separately for each query-target pair, without combining alignments to different reference sequences. Matches exceeding 95% nucleotide identity and 80% query coverage were retained. Each query was assigned to every reference resource containing a qualifying match, and intersections were summarized to distinguish sequences matching one or multiple resources.

### Predicted host taxonomy and food-category distributions

Putative hosts were assigned using HOTSPOT (30) with its publicly available pretrained models and reference databases. Predicted host lineages were summarized at the family level. Catalog-wide assignment rates were calculated using the 181,871 unique rPLATE PPCs as the denominator, counting exact sequence duplicates once. Host-family distributions across source categories were summarized separately using all 213,474 retained rPLATE PPC occurrences, without collapsing exact sequence duplicates. For each family, PPC occurrences were pooled within each source category, and each category’s contribution was calculated relative to the total number of occurrences assigned to that family across all categories.

### Functional potential profiling

Protein-coding sequences were predicted using Prodigal (31) in metagenomic mode, and small open reading frames were identified using SmORFinder (32) in metagenomic mode. Predicted protein and peptide sequences were used for functional annotation as described below. Contig and protein identifiers provided the links between functional annotations and their source sequences.

#### Antimicrobial resistance and additional stress-related functions

Predicted proteins were analyzed using AMRFinderPlus (33) with the NCBI AMRFinderPlus database and Resistance Gene Identifier (RGI) with the Comprehensive Antibiotic Resistance Database (CARD) (34). AMRFinderPlus was run with --plus to include additional stress-response and virulence-associated determinants alongside AMR genes. RGI was run in protein mode with DIAMOND (35).

AMRFinderPlus annotations were separated according to their reported type, subtype and class, so that AMR genes and additional functions, including heat and metal resistance, were summarized separately. RGI annotations were summarized using the reported drug classes and AMR gene families. Tool-specific results were retained because the two databases use different classification schemes.

For cross-tool comparisons, annotations were linked using sample/run, contig and protein identifiers. Protein-level comparisons were restricted to unique rPLATE PPCs, counting annotations on exact sequence duplicates once. Shared detection required both tools to annotate the same protein identifier; agreement on the exact gene identity or resistance class was assessed separately. For drug-class comparisons, combined labels were separated into their component classes, with each protein counted once per corresponding class. Antibiotic-focused sequencing-run summaries excluded AMRFinderPlus calls assigned to QUATERNARY AMMONIUM and RGI calls assigned exclusively to disinfecting agents and antiseptics; RGI calls combining antibiotic- and biocide-associated labels were retained. A sequencing-run record was considered positive by both tools when each tool reported at least one qualifying annotation, regardless of whether the annotations referred to the same protein.

#### Carbohydrate-active enzymes and gene clusters

Predicted proteins were annotated for carbohydrate-active enzymes (CAZymes) using dbCAN3 (36). The annotation workflow integrated DIAMOND, profile-HMM searches and dbCAN-sub. HMM-based matches were filtered using an E-value <10^□15^ and profile coverage >0.35. CAZyme assignments were recorded for each sample and summarized by enzyme class and family.

CAZyme gene clusters (CGCs) were identified using the CGC-Finder component of dbCAN. Candidate clusters were defined from the genomic proximity of CAZymes and associated signature genes, including transporters and transcription factors, using the default clustering settings. CGC annotations were treated as predictions of gene organization and functional potential.

*Protein families and KEGG orthologs*. Predicted proteins were annotated using eggNOG-mapper v2 (37) and eggNOG 5.0 (38) with DIAMOND search mode. Pfam (39) and KEGG Orthology (KO) (40) assignments were extracted and summarized in separate feature-by-sample count matrices.

#### CRISPR-Cas systems

PPCs were analyzed using CRISPRCasTyper (41). Predicted CRISPR arrays, cas genes and cas operons were extracted from the output, together with system-type and subtype assignments where available. CRISPR arrays and cas-associated loci were summarized separately to distinguish their detection from the identification of a complete CRISPR-Cas system.

#### Candidate bioactive peptides

Candidate anticancer peptides were identified by applying MetaPepticon (42) to the predicted peptide sequences using its default settings. Candidate antimicrobial peptides were identified using Macrel (43) in peptide-input mode with default settings. ACP and AMP classifications were reported as computational predictions of potential activity.

### Database accessibility

The PLATE website was developed as a browser-based interface for exploring food-associated PPCs and their annotations. Sample metadata and annotation outputs were organized using sample/run, contig and protein identifiers to link records across annotation datasets. The interface provides sample browsing and filtering, together with sample-specific pages displaying source accessions, provenance and associated annotation results. Functional annotation tables allow inspection of predicted gene content, while graphical CAZyme gene-cluster tracks display the organization of annotated genes within candidate clusters. Export functions support the retrieval of both sample metadata and sequences for categories selected by the user. PLATE is freely accessible at https://www.plate-db.com.

### Case study

A focused case study was performed to demonstrate PLATE’s utility. Quijada et al. (2025) reported the macrolide-associated gene pairs *msr(E)-mph(E)* and *msr(A)-mph(C)* on PPCs from food-production environments, particularly processing, ripening and packing rooms (44). Motivated by these findings, the distribution, same-contig co-occurrence and sequence conservation of selected macrolide-associated AMR genes across the food metagenomic datasets represented in PLATE were examined.

We searched “MACROLIDE” keyword in the PLATE website across all metadata and annotations and then downloaded sequence, metadata and annotation files for these results from the website.

AMRFinderPlus and RGI annotations were screened for PPCs carrying at least one exact MACROLIDE call and at least one exact MACROLIDE/STREPTOGRAMIN call on different protein identifiers. rPLATE retention was determined from contig identifiers in the detailed rPLATE AMRFinderPlus output. Within the retained rPLATE candidates, *msr(E)-mph(E), msr(G)-mef(F)*, and *msr(A)-mph(C)* were identified by requiring both exact AMRFinderPlus element symbols on the same contig. RGI support required corresponding Best_Hit_ARO identities at both AMRFinderPlus protein identifiers. Matching RGI calls were an additional annotation-support criterion and were not required for inclusion in the AMRFinderPlus-defined sequence-analysis set.

All rPLATE PPCs carrying mph(E) + msr(E) were compared pairwise using Smith-Waterman local alignment. Scores were +2 per match, □3 per mismatch and □5 per gap nucleotide under a linear gap model. Both target orientations were evaluated; the higher-scoring orientation was retained, with ties retaining the forward orientation. Identity was matches divided by alignment columns, including gaps. Coverage was the aligned coordinate span divided by full contig length. Minimum reciprocal coverage was the smaller of the two contig coverages. Shorter-contig coverage was the coverage of the physically shorter contig; the smaller coverage was used for equal-length PPCs. Cross-project alignments were summarized using thresholds of 80% minimum reciprocal coverage and 95% shorter-contig coverage.

Sensitivity analyses assessed exact-sequence redundancy and dependence on individual BioProjects. Whole-contig duplicates were identified allowing reverse-complement orientation, and summaries were recalculated after retaining one representative per identical sequence within each BioProject. Each BioProject was then omitted in turn to assess the persistence of the sequence-containment pattern.

## RESULTS

### Dataset coverage and analysis workflows

We assembled a global collection of 4,690 food-associated metagenomic datasets comprising accessions from cFMD (n = 3,990), NCBI (n = 645) and MGnify (n = 53), together with two samples sequenced in house (**Figure 1A**).

**Figure 1.**
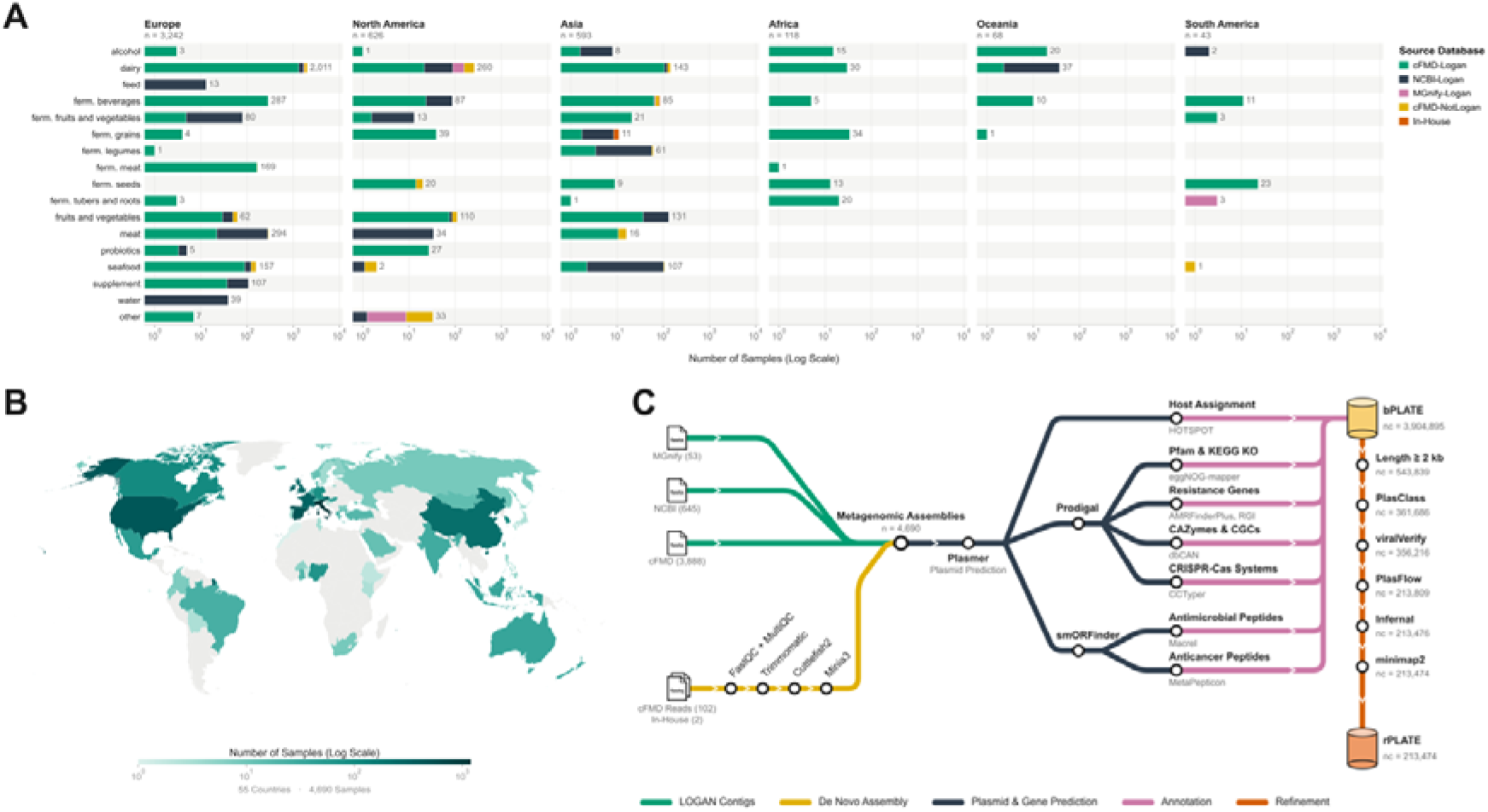
Dataset coverage and analysis workflow of the Plasmid Atlas of Food Ecosystems (PLATE). (A) Distribution of all included metagenomic datasets across the 17 source categories, faceted by continent. Stacked segments indicate dataset and assembly provenance, as defined in the legend. (B) Geographical distribution of food metagenome datasets. The color scale represents sample counts. (C) Overview of data collection, contig assembly or retrieval, plasmid prediction, catalog refinement, coding-sequence prediction, host assignment and functional annotation.

Preassembled contigs were obtained from Logan for 4,586 public datasets, whereas the remaining 102 public datasets and two in-house datasets were assembled from raw reads. Dataset sources, metadata, inclusion information and assembly provenance are given in **Supplementary Table S1**. The collection spans 55 countries and 17 source categories, encompassing fermented and non-fermented foods alongside feed, supplements and water. Dairy accounted for 2,481 datasets (52.9%), followed by fermented beverages (n = 485; 10.3%), meat (n = 344; 7.3%) and fruits and vegetables (n = 303; 6.5%). The geographical distribution of processed datasets is shown in **Figure 1B**. We applied two workflows with different filtering stringencies to generate the broad PLATE (bPLATE) and refined PLATE (rPLATE) catalogs (**Figure 1C**).

### Benchmarking plasmid prediction and refinement

We benchmarked the bPLATE and rPLATE workflows to evaluate their ability to distinguish plasmid-derived fragments from chromosomal sequences. The benchmarking dataset comprised plasmid fragments from PLSDB and chromosomal fragments from cFMD, with closely matched length distributions (**Figure 2A**). bPLATE identified 93.6% of plasmid fragments with a false positive rate (FPR) of 0.72%. As chromosomal sequences greatly outnumber plasmid sequences in food metagenomes, even a low FPR can yield a considerable proportion of false plasmid calls. By retaining only predictions supported by complementary classification approaches, rPLATE reduced the FPR more than threefold to 0.21% at the expense of lower recall (72.6%). The reduction in recall was most pronounced for short (2-5 kb) fragments (**Figure 2B**).

**Figure 2.**
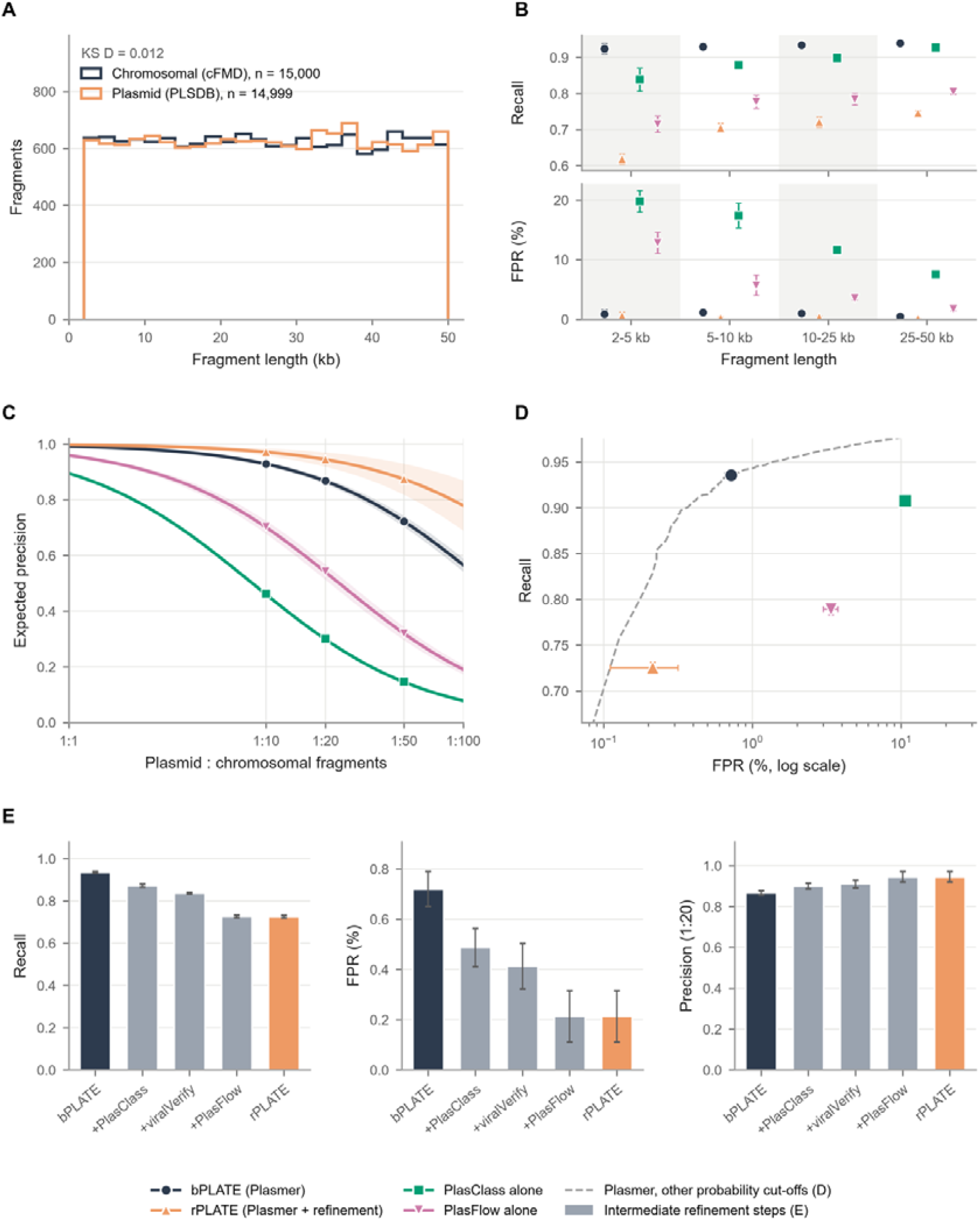
Benchmarking plasmid prediction workflows and the effects of sequential refinement. (A) Length distributions of plasmid fragments from PLSDB and chromosomal fragments from cFMD spanning 2-50 kb. (B) Recall and false positive rate (FPR) across four fragment-length intervals for bPLATE, rPLATE, and independently applied PlasClass and PlasFlow. (C) Expected precision across assumed plasmid-to-chromosomal fragment ratios, calculated from the benchmark recall and FPR. (D) Recall–FPR comparison, with FPR shown on a logarithmic scale. The dashed grey curve indicates Plasmer performance at alternative probability cut-offs. (E) Changes in recall, FPR and expected precision at an assumed plasmid-to-chromosomal fragment ratio of 1:20 during sequential refinement. Navy circles represent bPLATE, orange triangles represent rPLATE, green squares represent PlasClass applied independently, and pink inverted triangles represent PlasFlow applied independently in panels B-D. Grey bars indicate intermediate refinement stages in panel E.

The precision advantage of refinement increased as the assumed proportion of plasmid-derived fragments decreased; at a plasmid-to-chromosomal fragment ratio of 1:50, expected precision increased from 0.72 for bPLATE to 0.87 for rPLATE (**Figure 2C**). When applied independently to all fragments, PlasClass and PlasFlow showed FPRs of 10.6% and 3.4%, respectively (**Figure 2D**), whereas their sequential application to Plasmer predictions substantially reduced false positive calls, highlighting the benefit of combining complementary prediction methods. Within the refinement chain, PlasFlow produced the largest decrease in recall (**Figure 2E**). FPRs remained below 2% for bPLATE and below 1% for rPLATE across the source categories represented in the benchmark. Accordingly, bPLATE is suited to sensitive, exploratory detection, whereas rPLATE provides a more stringently filtered catalog with lower false-positive rates in the benchmark. Detailed performance metrics are provided in **Supplementary Table S2**.

### PLATE composition and comparison with reference plasmid databases

bPLATE comprises 3,904,895 putative plasmid contigs (PPCs) from 4,635 datasets, while rPLATE comprises 213,474 PPCs, 181,871 unique PPCs and 63,820 PPC clusters from 4,349 datasets (**Figure 3A**). Sequential filtering outcomes, including human-reference and rRNA exclusions, are detailed in **Supplementary Table S3**. Mean PPC length per accession ranged from about 0.5 kb to 7 kb in bPLATE (**Supplementary Figure S1A**), with both extremes in dairy, and from about 2 kb to 19 kb in rPLATE, with both extremes in fruits and vegetables (**Supplementary Figure S1B**).

**Figure 3.**
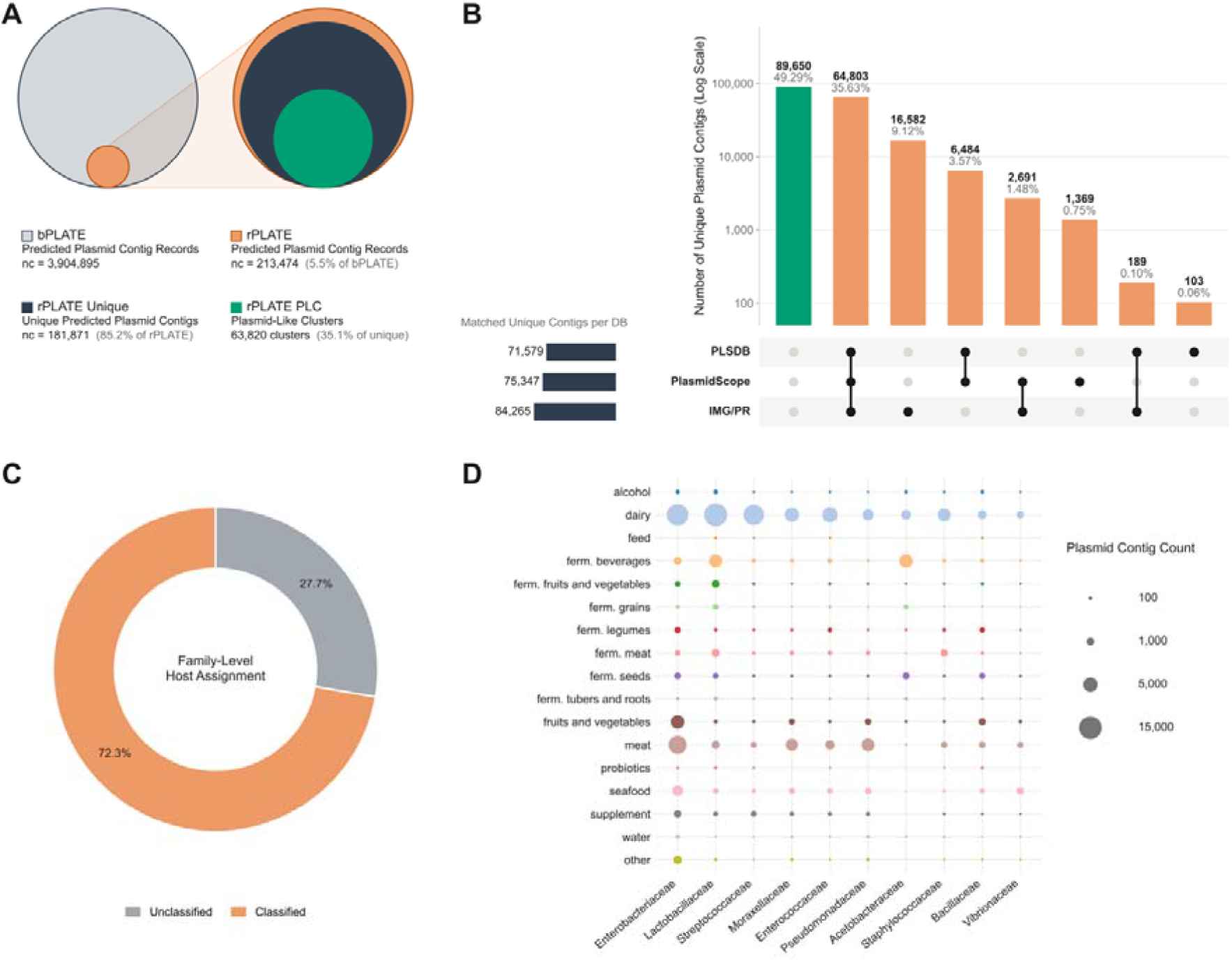
Catalog refinement, reference representation and predicted hosts of PPCs. (A) Sequence counts and membership of bPLATE and rPLATE, and sequence redundancy within rPLATE. (B) UpSet plot showing intersections among unique rPLATE PPCs with qualifying matches to the indicated reference plasmid databases. Intersection bars count PLATE PPCs matching the corresponding combination of databases. (C) Proportions of the 181,871 unique rPLATE PPCs assigned and unassigned to predicted hosts at the family level using HOTSPOT. (D) Pooled counts of retained rPLATE PPC occurrences by predicted host family and source category. Bubble size represents the number of PPC occurrences; exact sequence duplicates are counted separately, and counts are not normalized by sample number.

Existing plasmid resources contain a varying number of sequences explicitly labelled as food-associated (**Table 1**). To assess reference representation beyond food-source labelling, we compared all unique rPLATE PPCs with all sequences in PLSDB (12), PlasmidScope (14) and IMG/PR (13), regardless of their source annotations. Using thresholds of >95% nucleotide identity and >80% query coverage, 92,221 rPLATE PPCs matched at least one reference resource. Of these, 74,167 matched multiple databases and 18,054 matched only one, while the remaining 89,650 rPLATE PPCs lacked a qualifying match (**Figure 3B**). Detailed reference matches and database assignments are provided in **Supplementary Table S4**.

**Table 1.**
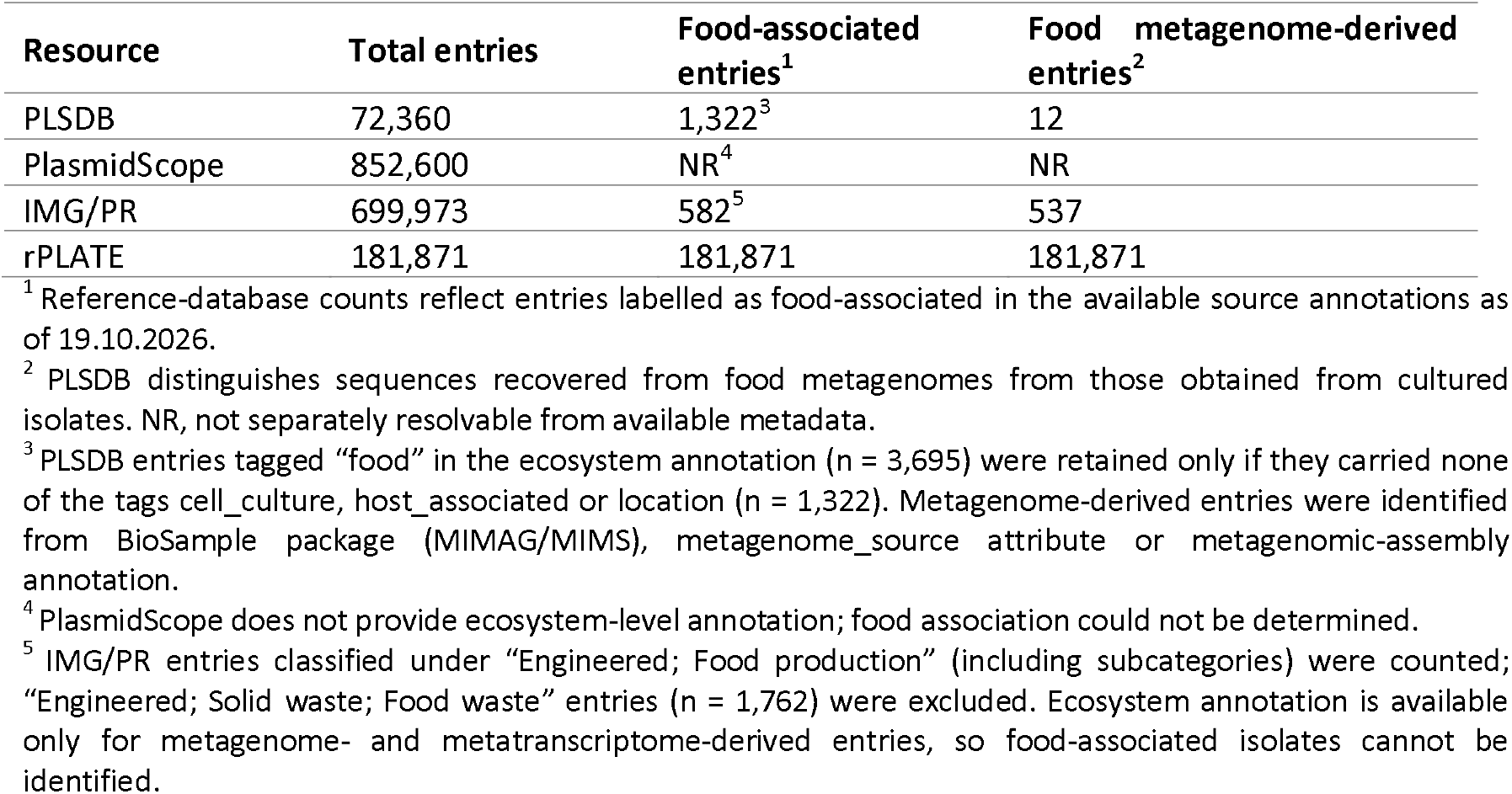
Unique PPC counts in rPLATE and food-associated entries in existing plasmid databases.

### Taxonomic and functional annotations

#### Predicted hosts and their distribution across source categories

Among the 181,871 unique rPLATE PPCs, HOTSPOT assigned 131,459 (72.3%) at the family level (**Figure 3C**). Detailed host assignments are provided in *Supplementary Table S5*.

To describe host-family distributions across source categories, we counted all retained rPLATE PPC occurrences, including repeated occurrences of identical sequences across source records. The most frequently assigned families were *Enterobacteriaceae* (35,363 occurrences), *Lactobacillaceae* (26,010) and *Streptococcaceae* (13,989). Dairy contributed 60.7% of *Lactobacillaceae* and 84.7% of *Streptococcaceae* occurrences, whereas 62.0% of *Enterobacteriaceae* occurrences were recovered outside dairy, with substantial contributions from meat and fruits and vegetables. Fermented beverages contributed 56.3% of *Acetobacteraceae* occurrences, while meat was the largest source category for *Pseudomonadaceae* (44.3%) (**Figure 3D**).

#### Antibiotic, biocide and heat-resistance annotations

AMRFinderPlus identified 2,153 AMR-associated proteins on 1,694 rPLATE PPCs from 987 sequencing-run records. The most frequent reported class labels were aminoglycoside (654), tetracycline (630) and beta-lactam (355), together accounting for 76.1% of AMRFinderPlus AMR detections. Dairy contributed 872 annotations (40.5%), followed by meat (644; 29.9%) and fruits and vegetables (177; 8.2%) (**Figure 4A**).

**Figure 4.**
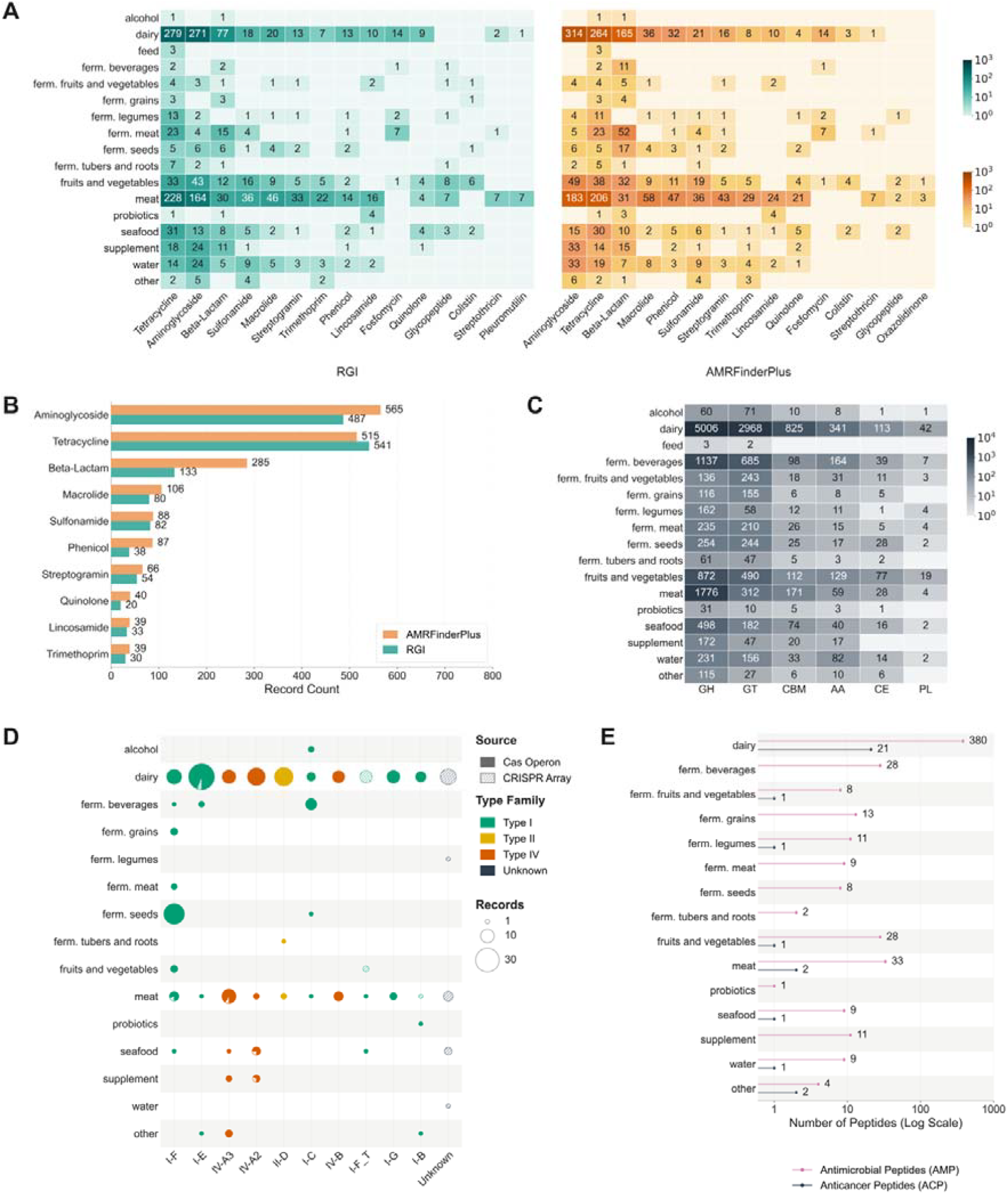
Functional annotations of PPCs in rPLATE. (A) Heatmaps showing protein-level annotation counts for the displayed antibiotic-resistance classes across source categories, based on AMRFinderPlus and RGI. Counts include repeated occurrences on identical sequences recovered from different records. Proteins with combined class labels contribute once to each corresponding displayed class. Biocide classes are omitted. (B) Numbers of resistance-gene records per drug class detected by AMRFinderPlus and RGI, for the ten classes with the most AMRFinderPlus records. Counts are restricted to unique rPLATE PPCs, and records carrying combined class labels are counted once under each component class. (C) Distribution of carbohydrate-active enzyme (CAZyme) classes across source categories. (D) Distribution of CRISPR-Cas annotations across source categories. (E) Lollipop plot showing numbers of predicted antimicrobial peptides (AMPs) and anticancer peptides (ACPs) across source categories.

RGI identified 2,308 resistance-associated proteins on 1,773 rPLATE PPCs from 1,019 sequencing-run records. Meat contributed the largest number of RGI annotations (906), followed by dairy (754), fruits and vegetables (161) and seafood (130). The most frequent single-class labels were tetracycline antibiotic (657), aminoglycoside antibiotic (556) and disinfecting agents and antiseptics (553). The latter 553 records contained only the biocide-associated class label, whereas six additional records combined antibiotic- and biocide-associated labels. The most frequently reported individual gene was *qacJ*, with 411 detections.

Restricting annotations to unique rPLATE PPCs retained 1,816 AMRFinderPlus AMR-associated protein records (84.3% of all rPLATE detections) and 1,860 RGI resistance-associated protein records (80.6%). Matching protein identifiers between the tools identified 1,393 shared detections, representing 76.7% and 74.9% of the respective protein sets. Across all rPLATE PPC occurrences, including repeated occurrences of identical sequences, the corresponding counts were 1,653 shared, 500 AMRFinderPlus-only and 655 RGI-only detections.

Drug-class comparisons were summarized separately by assigning proteins with combined class labels to each corresponding component class. On unique rPLATE PPCs, AMRFinderPlus and RGI returned comparable numbers of class assignments for tetracycline (515 and 541), aminoglycoside (565 and 487) and sulfonamide (88 and 82), whereas AMRFinderPlus returned more beta-lactam (285 versus 133) and phenicol (87 versus 38) assignments (**Figure 4B**). Detailed protein-level annotations are provided in **Supplementary Table S6**.

Comparison with the bPLATE AMRFinderPlus annotation export showed that 2,153 of 10,609 AMR-associated protein records (20.3%) were retained in rPLATE PPC occurrences. Aminoglycoside and tetracycline annotations remained prominent after refinement, but the raw category ranking changed from meat followed by dairy in bPLATE to dairy followed by meat in rPLATE. Changes in class composition are shown in **Supplementary Figure S2A**. Under the antibiotic-focused filtering criteria, which excluded biocide-only detections, 986 rPLATE sequencing-run records were positive by AMRFinderPlus, 874 by RGI and 845 by both tools (**Supplementary Figure S2B**).

AMRFinderPlus additionally identified 221 heat-resistance-associated proteins on 93 rPLATE PPCs, compared with 2,674 protein records in the bPLATE export. Dairy accounted for 160 rPLATE heat-resistance annotations (72.4%), followed by meat (17), fruits and vegetables (14) and supplements (14). Heat-resistance profiles are shown separately in **Supplementary Figure S3**.

#### Carbohydrate-active enzymes and predicted gene clusters

CAZyme annotation identified 19,221 proteins on 15,959 rPLATE PPCs. Glycoside hydrolases (GH; 10,865 proteins) and glycosyltransferases (GT; 5,907) were the most strongly represented classes and were the two largest classes across all 17 source categories. GH predominated in most categories, whereas GT exceeded GH in alcohol, fermented fruits and vegetables, and fermented grains. Dairy contributed the largest share of CAZyme-annotated proteins (47.6%), followed by meat and fermented beverages (**Figure 4C**).

CGC prediction identified 3,685 candidate clusters on 3,666 rPLATE PPCs. All clusters contained CAZymes and 95.1% (3,503) also contained predicted transporter annotations, with some additionally containing signal-transduction proteins, transcription factors, peptidases or sulfatases. Cluster architecture varied in gene content, size and orientation, encompassing 2-16 genes over spans of 662–19,226 bp. Most clusters contained genes on a single strand (79.6%), whereas 20.4% included genes on both strands. CAZyme annotations, cluster coordinates, gene identities and orientations are provided in **Supplementary Table S7**.

#### KEGG orthologs and Pfam domains

eggNOG-mapper assigned 270,228 KEGG Orthology (KO) records spanning 6,550 KOs and 890,008 Pfam records spanning 6,851 domains to rPLATE proteins (Supplementary Table S8). The 15 most frequent KOs accounted for 16.8% of all KO records, whereas the 15 most frequent Pfam domains accounted for 15.1% of Pfam records (Supplementary Figure S4).

The most frequent KOs primarily represented plasmid maintenance, DNA transfer and DNA processing functions. These included the partitioning proteins ParA and ParB, the type IV secretion system coupling protein VirD4, and the conjugal transfer ATPase TraC, alongside DNA topoisomerase III, DNA polymerase V (UmuC), and single-stranded DNA-binding protein. Four KOs associated with DNA methylation and restriction-modification systems were also among the top 15, while two transposase KOs together accounted for 4,752 annotation records. Transport-related functions included an ABC-2 transporter ATPase and a zinc/cadmium-exporting ATPase.

The relative representation of these functions varied across source categories (**Supplementary Figure S4**). ParA was the most frequent KO in most categories, while VirD4 predominated in fermented meat and ParB in seafood and water. VirD4 accounted for 14.3% of top-15 KO records in dairy and 6.2% in fruits and vegetables, whereas TraC contributed 8.7% in seafood and 2.1% in dairy. ParB represented 21.8% of top-15 KO records in water, compared with 8.2% in dairy.

#### CRISPR-Cas systems

CRISPRCasTyper reported CRISPR arrays on 43 PPCs and Cas-associated loci on 227 PPCs in rPLATE. Cas-associated loci represented types I-V, with I-F (49 records), I-E (38) and IV-A3 (26) most frequently reported. Dairy accounted for the largest share of both CRISPR-array records (58.1%) and Cas-associated locus records (57.3%), followed by meat (*Figure 4D*). Source records, subtype assignments and reported scores are provided in *Supplementary Table S9*.

#### Antimicrobial and anticancer peptides

Peptide screening identified 554 candidate antimicrobial peptides and 30 candidate anticancer peptides, corresponding to 186 and 16 unique amino-acid sequences, respectively. Dairy contributed 380 AMP candidates (68.6%) and 21 ACP candidates (70.0%) (**Figure 4E**). Of the AMP candidates, 272 were predicted to be non-hemolytic and 282 hemolytic; all 30 ACP candidates were predicted to be non-toxic. Peptide sequences, source records and prediction outputs are provided in **Supplementary Table S10**.

#### Web interface and data access

The PLATE web interface provides access to sample metadata, PPCs and their associated annotations at https://www.plate-db.com (**Figure 5A**). PLATE website supports browsing, filtering and downloading of sample records and annotations (**Figure 5B**). Selecting a sample opens a dedicated page displaying source accessions, provenance and annotation results linked through sample/run, contig and protein identifiers (**Figure 5C**). The annotations on PPCs belonging to refined catalog are marked with a darker color (**Figure 5D**). Functional annotation tables in the PLATE website support inspection of predicted gene content, while graphical CAZyme gene-cluster tracks display annotated gene organization. Export functions enable retrieval of sample metadata and sequences for further analysis.

**Figure 5.**
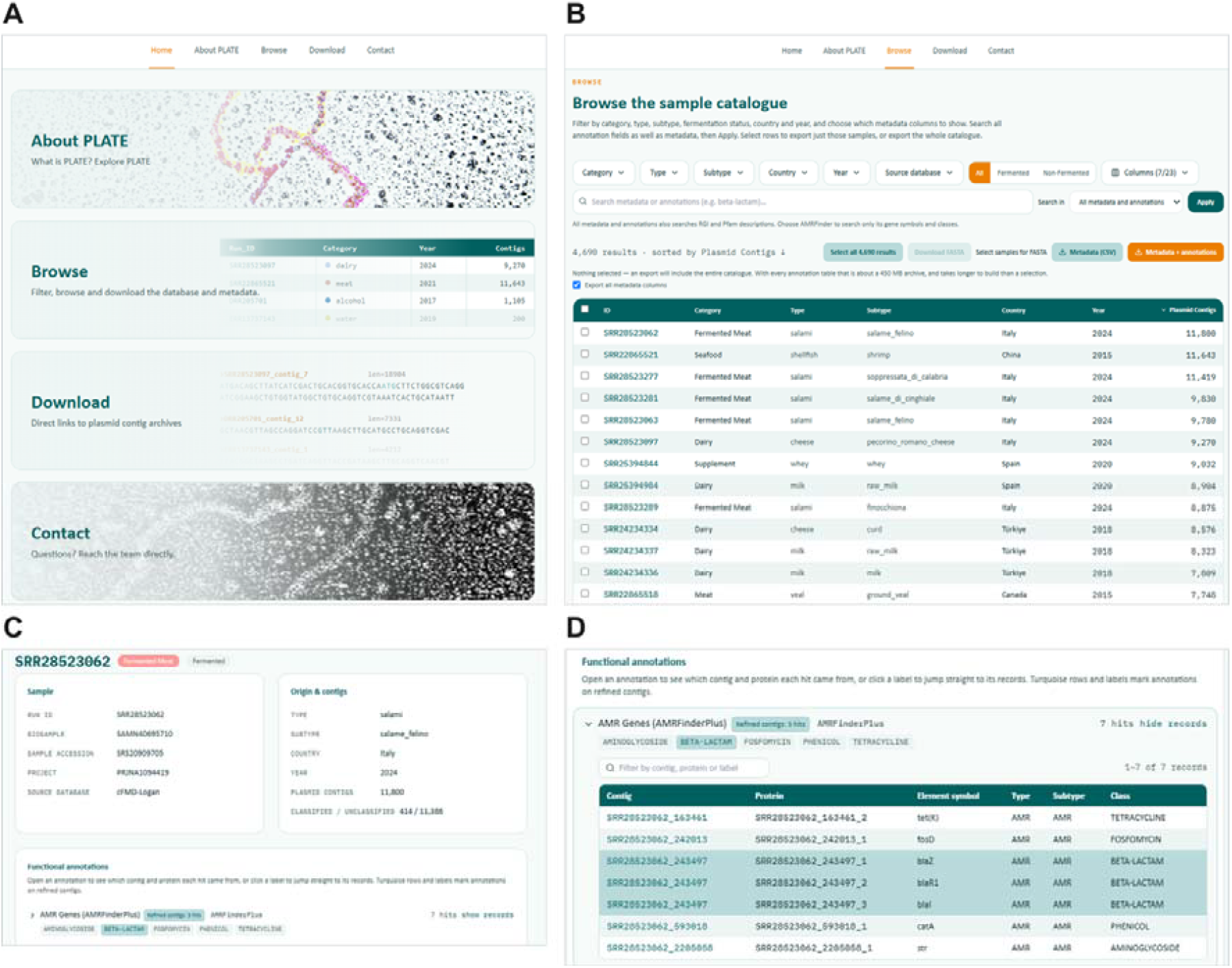
The PLATE web interface. (A) The home page links to the main sections: About PLATE, Browse, Download and Contact. (B) The Browse page shows the sample catalogue (4,690 sequencing runs). Datasets can be filtered by category, type, subtype, country, year, source database and fermentation status, and the columns shown can be chosen. A search box queries metadata and functional annotations together or separately. Results can be sorted (here by plasmid contig count), and users can export PPC sequences (FASTA) for selected datasets, metadata alone (CSV), or metadata with annotations. (C) The record page for each dataset shows the sample identifiers (run, BioSample, sample accession and BioProject), the source database, the sample’s origin, its number of PPCs, and how many of these are classified or unclassified. Below these, the sample’s functional annotations are summarized. (D) The functional annotations are summarized for each sample record, listing each hit with its contig, protein, element symbol, type, subtype and resistance class, and the class labels can be used to filter the table. Turquoise rows and labels mark hits on rPLATE PPCs.

### Case study: macrolide-associated gene co-occurrence and sequence conservation

Screening bPLATE for PPCs carrying both MACROLIDE and MACROLIDE/STREPTOGRAMIN annotations on separate proteins identified 63 candidate PPCs from 57 sequencing-run records by AMRFinderPlus. These spanned 17 BioProjects, 12 countries and seven source categories, including water. Of these, 37 PPCs from 35 records were retained in rPLATE, representing 11 BioProjects, nine countries and six source categories. Applying the corresponding exact RGI drug classes identified 45 bPLATE PPCs, of which 29 were retained in rPLATE. All RGI-defined candidates were also among the AMRFinderPlus-defined candidates (**Figure 6A**). Candidate identifiers and source metadata are provided in **Supplementary Table S11**.

**Figure 6.**
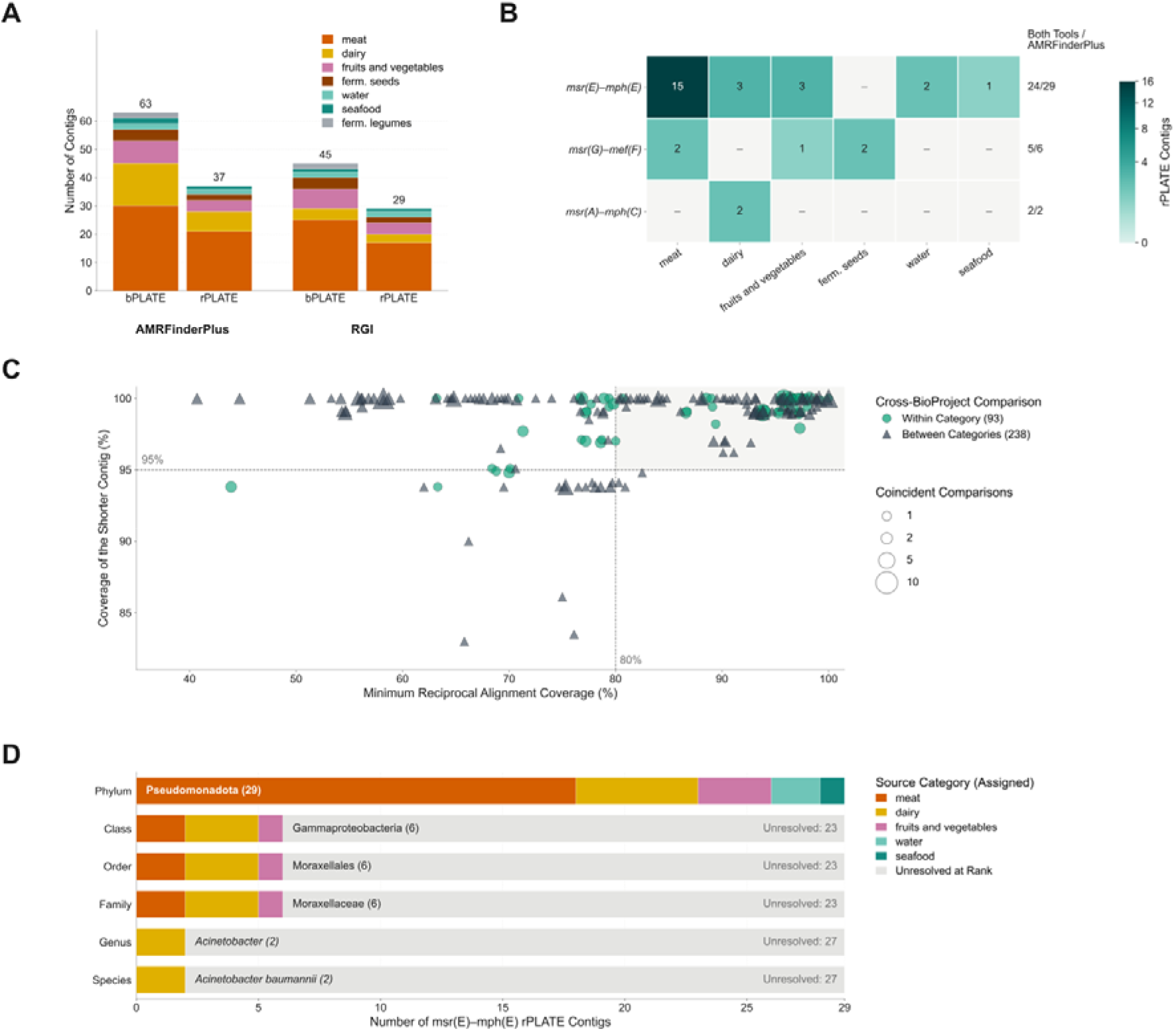
Distribution and sequence context of recurrent macrolide-associated gene combinations in PLATE. (A) Numbers of bPLATE and rPLATE PPCs carrying separate proteins annotated under the exact resistance classes macrolide and macrolide/streptogramin by AMRFinderPlus and RGI. Bars are colored by source category. (B) Heatmap showing the numbers of rPLATE PPCs on which both AMRFinderPlus and RGI identified both genes of *msr(E)-mph(E), msr(G)-mef(F)* or *msr(A)-mph(C)*, across source categories. Gene-pair occurrence requires both genes to be detected on the same PPC. Values at right give PPCs supported by both tools out of AMRFinderPlus-defined PPCs. (C) Minimum reciprocal alignment coverage versus coverage of the shorter PPC for cross-BioProject comparisons of the 29 *msr(E)-mph(E)*-carrying rPLATE PPCs. Colors and shapes distinguish within-category and between-category comparisons; marker size indicates the number of coincident comparisons. Reference lines mark 80% minimum reciprocal coverage and 95% shorter-contig coverage; the shaded area marks comparisons meeting both thresholds. (D) Predicted host taxonomy of the 29 *msr(E)-mph(E)-*carrying PPCs from phylum to species. Colored segments show PPCs assigned at each rank by source category; grey segments show PPCs unresolved at that rank.

AMRFinderPlus gene symbols identified *msr(E)-mph(E)* on 29 retained PPCs, *msr(G)-mef(F)* on six and *msr(A)-mph(C)* on two. Matching RGI identities supported both genes on 24, five and two PPCs, respectively, giving 31 PPCs with both gene identities supported by both tools (**Figure 6B**). *msr(E)-mph(E)* was the most widely distributed combination, detected in meat, dairy, fruits and vegetables, water and seafood.

Sequence characterization included all 29 AMRFinderPlus-defined *msr(E)-mph(E)* carrying PPCs (2,010-5,326 bp), originating from 29 metadata records across eight BioProjects, seven countries and five source categories. Pairwise comparison of these sequences yielded 406 alignments, including 331 between different BioProjects. Cross-project alignments showed 99.80-100% nucleotide identity across aligned regions. Of these, 186 covered at least 80% of both PPCs, 299 covered at least 95% of the shorter PPC, and 183 (55.3%) met both thresholds (**Figure 6C**). Comparisons meeting both thresholds linked 19 of the 28 possible BioProject pairs and included 123 of 238 between-category and 60 of 93 within-category comparisons. Among the 145 comparisons below 80% minimum reciprocal coverage, 116 nevertheless covered at least 95% of the shorter PPC. Thus, most comparisons with low reciprocal coverage involved extensive containment of a shorter sequence within a longer one. The 29 PPCs represented 26 exact whole-sequence types, with three duplicate groups confined to individual BioProjects. After deduplication, 240 of 268 cross-project comparisons (89.6%) covered at least 95% of the shorter PPC, and 156 (58.2%) met both thresholds. Omitting each BioProject in turn from the deduplicated set retained the shorter-contig coverage threshold in 86.1-97.9% of the remaining comparisons. In every case, all remaining BioProjects stayed linked by at least one comparison meeting both thresholds.

All 29 msr(E)-mph(E) PPCs were assigned to *Pseudomonadota*, but only six were resolved below phylum. All six belonged to *Gammaproteobacteria, Moraxellales* and *Moraxellaceae* and came from dairy, meat and fruit-and-vegetable datasets. Two kefir PPCs from the same BioProject were further assigned to *Acinetobacter baumannii* and were exact duplicates (**Figure 6D**). These findings indicate recurrence of highly conserved resistance-associated regions within predominantly shared or nested assembled sequences across source records, consistent with a *Moraxellaceae* host where taxonomy could be resolved.

## DISCUSSION

PLATE connects putative plasmid contigs (PPC) recovered from food-associated metagenomes with functional annotations and curated source metadata, enabling their gene content to be examined within its sequence and food-source context. Its contribution complements established plasmid resources, which provide reference sequences, annotations and tools for investigating plasmids across diverse environments. PLATE organizes food-associated candidates into two connected catalogs and links them to retrievable sequences and source records. This framework supports analyses ranging from catalog-wide annotation profiles to the genomic context of recurrent gene combinations, as illustrated by the macrolide-associated resistance case study.

Comparison with existing plasmid resources identified qualifying matches for 92,221 rPLATE PPCs, whereas 89,650 lacked a match under the applied identity and query-coverage criteria. These results indicate that PLATE contains both sequences represented in established resources and candidates whose representation remains unresolved at the selected thresholds. The unmatched fraction should not be interpreted directly as novel plasmid diversity, because sequence divergence, incomplete reference coverage, assembly fragmentation and classification errors can all contribute to the absence of qualifying matches. Likewise, extensive coverage of a PLATE PPC by a reference alignment does not establish recovery of the complete reference plasmid. However, the comparison provides a basis for prioritizing candidates for further characterization and assessing how food-derived sequences are represented in broader plasmid collections.

PLATE provides candidates for investigating antimicrobial resistance, microbial defense, carbohydrate utilization, and bioactive peptides. AMR annotations enable examination of resistance-gene distributions and same-contig co-occurrence across food sources, supporting candidate selection for further sequence-context and functional analyses. Predicted CAZyme gene clusters add information about local organization by linking CAZymes with transporters and other associated genes. CRISPR arrays and Cas-associated loci provide candidates for examining defense-related functions, while peptide screening identifies candidate sequences with potential antimicrobial or anticancer activity. Together, these annotations help prioritize candidates for experimental investigation of plasmid-associated functions.

It should be noted that PLATE’s datasets are unevenly distributed across food categories, and differences in sampling, sequencing depth and sample-processing and assembly procedures affect sequence and annotation recovery. Comparisons among food categories therefore require appropriate normalization and consideration of study-level dependence. The contig-based catalog may also contain fragmented or recurrent sequences, so entry counts do not represent distinct plasmids. Plasmid origin, host associations and functional annotations are computational predictions requiring further validation. Long-read sequencing, direct host-plasmid linkage and targeted functional assays could strengthen these assignments and clarify their biological relevance.

PLATE makes food-associated PPCs and their annotations accessible for comparative analysis and experimental follow-up. Its sequence-based case study demonstrates how users can move from resistance-class summaries to conserved gene regions and shared contig contexts. The resource’s value lies in making these findings traceable and reusable while exposing the effects of catalog selection and the uncertainty inherent in computational predictions.

## ACKNOWLEDGEMENTS

The authors thank Ali Eser for his support in website development. The numerical calculations reported in this paper were partially performed at TUBITAK ULAKBIM, High Performance and Grid Computing Center (TRUBA resources).

## AUTHOR CONTRIBUTIONS

Mehmet Umur Aslan: Data curation, Formal analysis, Investigation, Methodology, Software, Visualization, Writing—original draft, Writing—review & editing. Rıza Arda Kayacan: Data curation, Formal analysis, Investigation, Methodology, Software, Visualization, Writing—original draft. Süleyman Arif Söğütdelen: Data curation, Formal analysis, Investigation, Methodology, Software, Visualization, Writing—original draft. Özlem Sağıroğlu: Methodology, Validation, Writing—review & editing. Muzaffer Arıkan: Conceptualization, Funding acquisition, Supervision, Data curation, Investigation, Methodology, Project administration, Resources, Software, Validation, Writing— review & editing.

## SUPPLEMENTARY DATA

Supplementary material accompanies this preprint.

## DATA AVAILABILITY

The PLATE database is freely accessible at https://www.plate-db.com. All code used to construct PLATE and detailed documentation is available at https://github.com/arikanlab/PLATE. The broad catalog (bPLATE) and refined catalog (rPLATE) and their associated metadata are also available through Zenodo (https://doi.org/10.5281/zenodo.23691515). Raw shotgun metagenomic sequencing reads from the in-house boza and kefir samples have been deposited in the NCBI Sequence Read Archive under BioProject accession PRJNA1555283.

