## Supplementary Figure 1 for "PLATE: a global atlas of food metaplasmidomes"

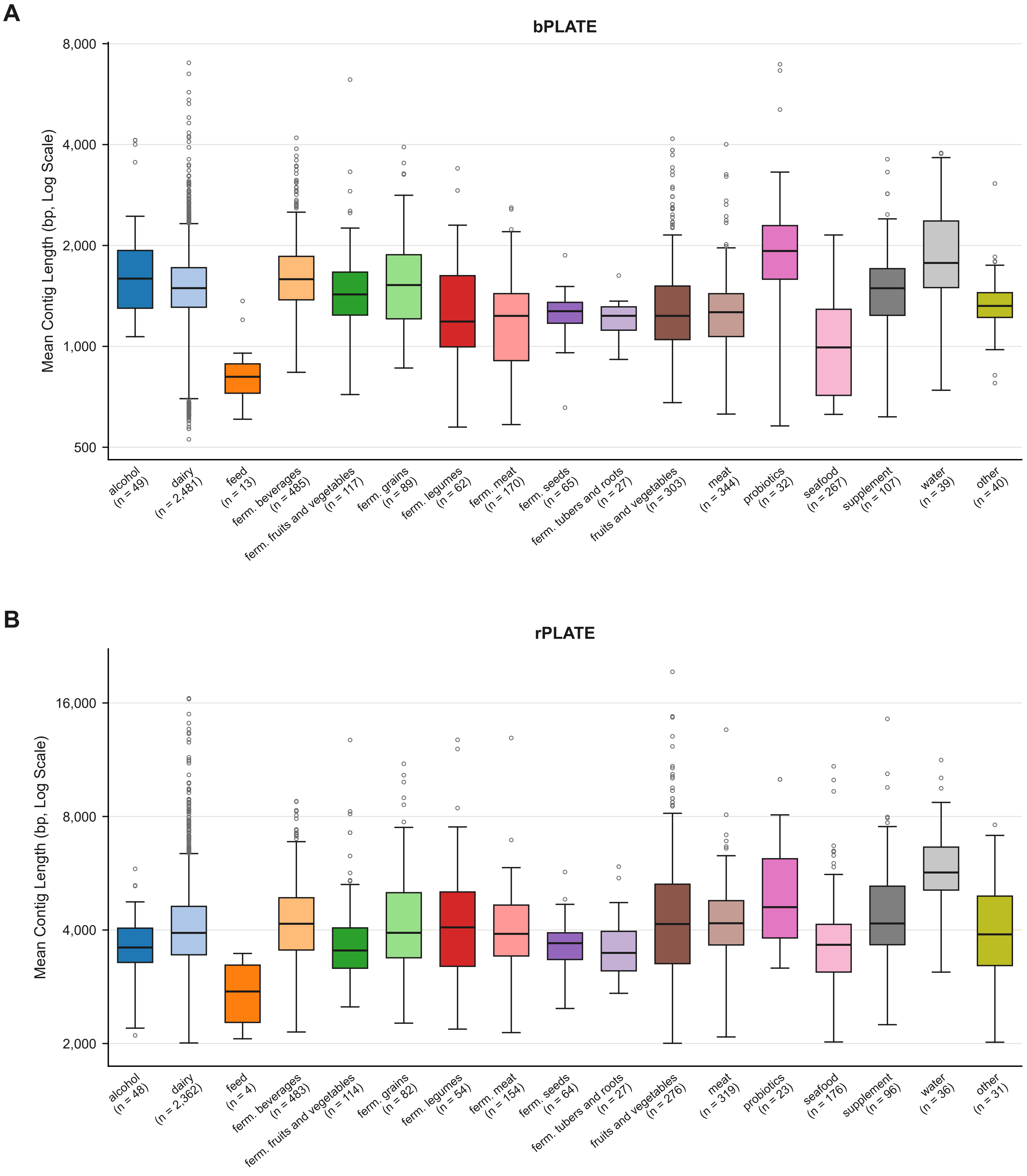
A
bPLATE
Mean Contig Length (bp, Log Scale)
8,000
4,000
2,000
1,000
500
alcohol (n = 49)
dairy (n = 2,481)
feed (n = 13)
ferm. beverages (n = 485)
ferm. fruits and vegetables (n = 117)
ferm. grains (n = 89)
ferm. legumes (n = 62)
ferm. meat (n = 170)
ferm. seeds (n = 65)
ferm. tubers and roots (n = 27)
fruits and vegetables (n = 303)
meat (n = 344)
probiotics (n = 32)
seafood (n = 267)
supplement (n = 107)
water (n = 39)
other (n = 40)
B
rPLATE
Mean Contig Length (bp, Log Scale)
16,000
8,000
4,000
2,000
alcohol (n = 48)
dairy (n = 2,362)
feed (n = 4)
ferm. beverages (n = 483)
ferm. fruits and vegetables (n = 114)
ferm. grains (n = 82)
ferm. legumes (n = 54)
ferm. meat (n = 154)
ferm. seeds (n = 64)
ferm. tubers and roots (n = 27)
fruits and vegetables (n = 276)
meat (n = 319)
probiotics (n = 23)
seafood (n = 176)
supplement (n = 96)
water (n = 36)
other (n = 31)
