## Supplementary Figure 2 for "PLATE: a global atlas of food metaplasmidomes"

A

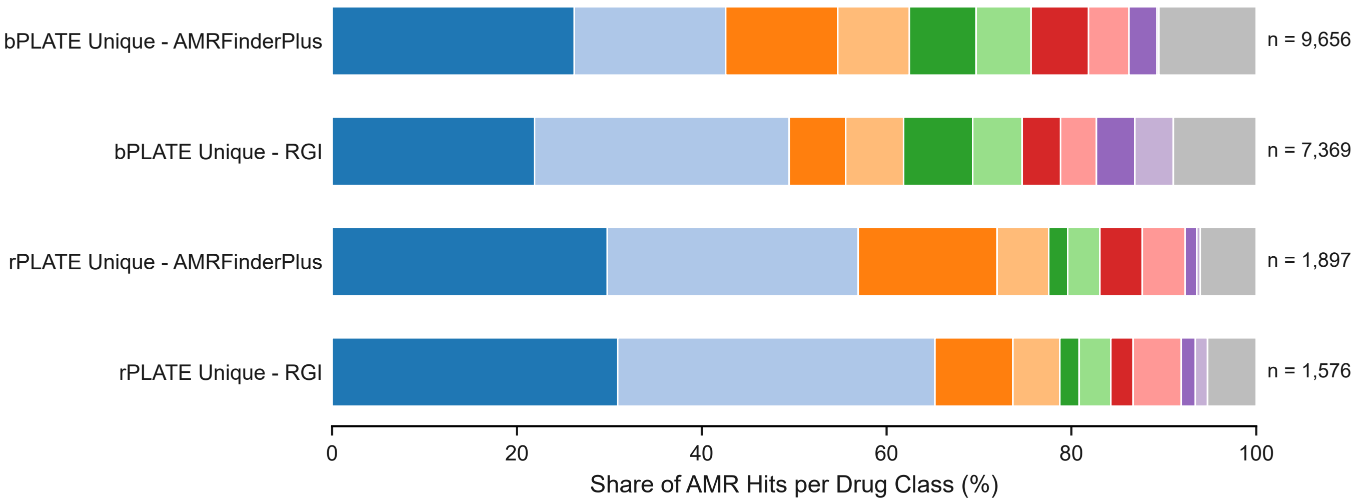
bPLATE Unique - AMRFinderPlus
bPLATE Unique - RGI
rPLATE Unique - AMRFinderPlus
rPLATE Unique - RGI
n = 9,656
n = 7,369
n = 1,897
n = 1,576
0
20
40
60
80
100
Share of AMR Hits per Drug Class (%)
Aminoglycoside
Tetracycline
Beta-Lactam
Macrolide
Lincosamide
Streptogramin
Phenicol
Sulfonamide
Fosfomycin
Glycopeptide
Other Classes

B

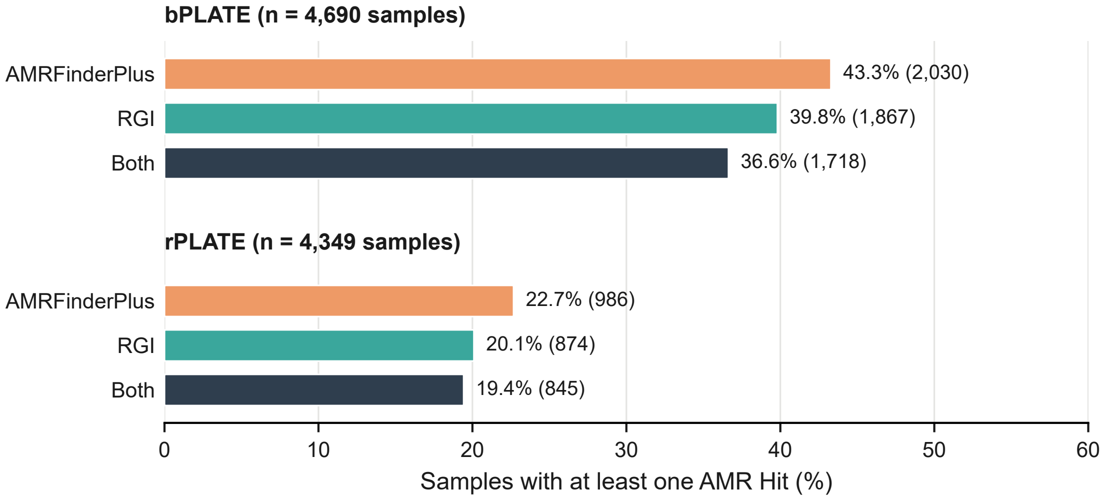
bPLATE (n = 4,690 samples)
AMRFinderPlus
RGI
Both
43.3% (2,030)
39.8% (1,867)
36.6% (1,718)
rPLATE (n = 4,349 samples)
AMRFinderPlus
RGI
Both
22.7% (986)
20.1% (874)
19.4% (845)
0
10
20
30
40
50
60
Samples with at least one AMR Hit (%)
