## Supplementary Figure 3 for "PLATE: a global atlas of food metaplasmidomes"

A

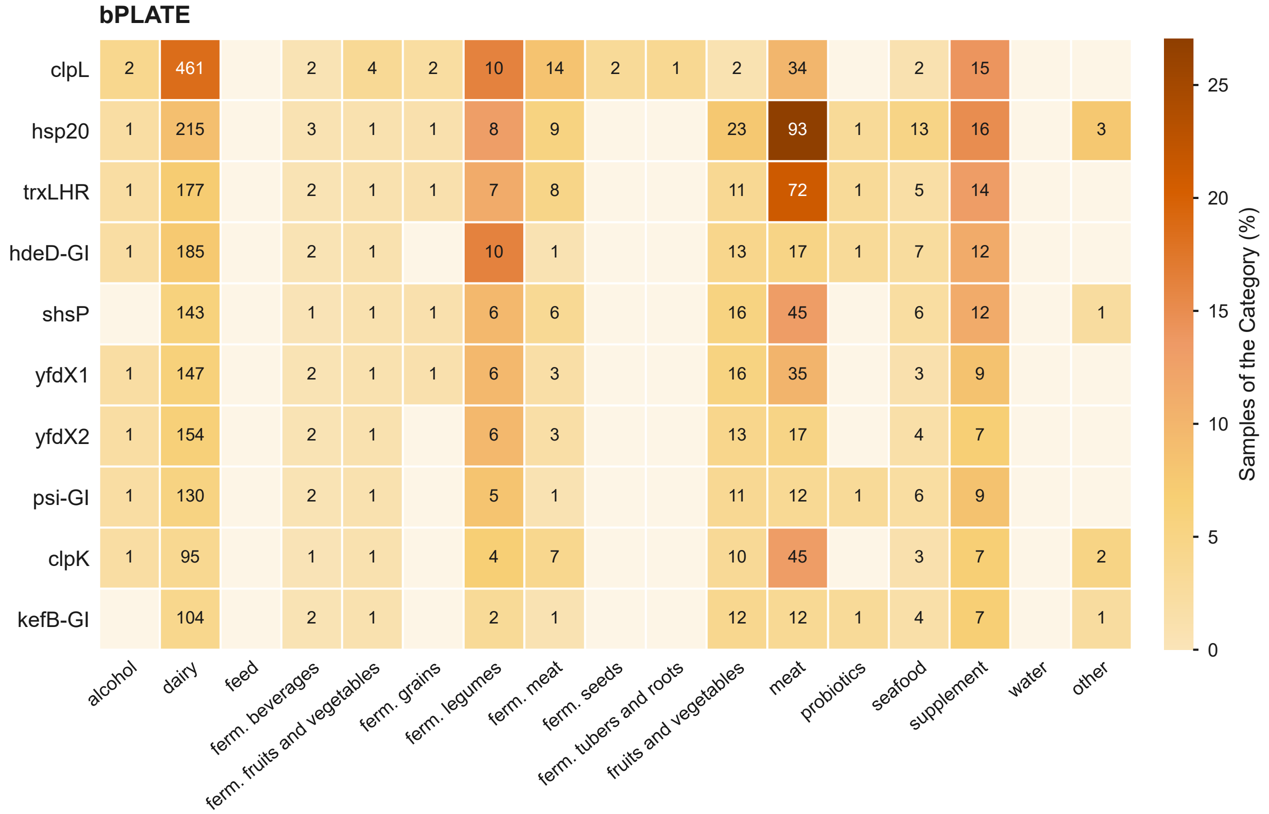
bPLATE
clpL
hsp20
trxLHR
hdeD-GI
shsP
yfdX1
yfdX2
psi-GI
clpK
kefB-GI
alcohol
dairy
feed
ferm. beverages
ferm. fruits and vegetables
ferm. grains
ferm. legumes
ferm. meat
ferm. seeds
ferm. tubers and roots
fruits and vegetables
meat
probiotics
seafood
supplement
water
other
Samples of the Category (%)
0
5
10
15
20
25

B

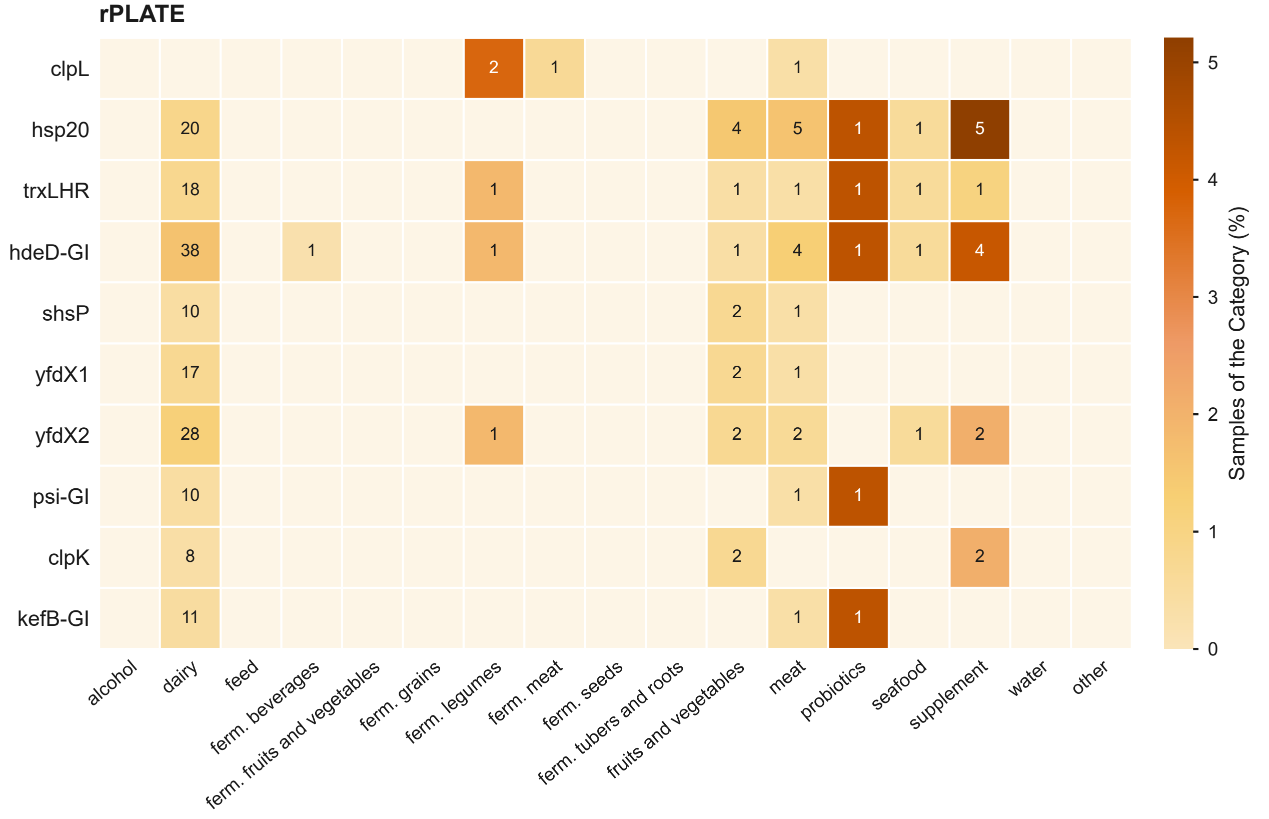
rPLATE
clpL
hsp20
trxLHR
hdeD-GI
shsP
yfdX1
yfdX2
psi-GI
clpK
kefB-GI
alcohol
dairy
feed
ferm. beverages
ferm. fruits and vegetables
ferm. grains
ferm. legumes
ferm. meat
ferm. seeds
ferm. tubers and roots
fruits and vegetables
meat
probiotics
seafood
supplement
water
other
Samples of the Category (%)
0
1
2
3
4
5
