## Supplementary Figure 4 for "PLATE: a global atlas of food metaplasmidomes"

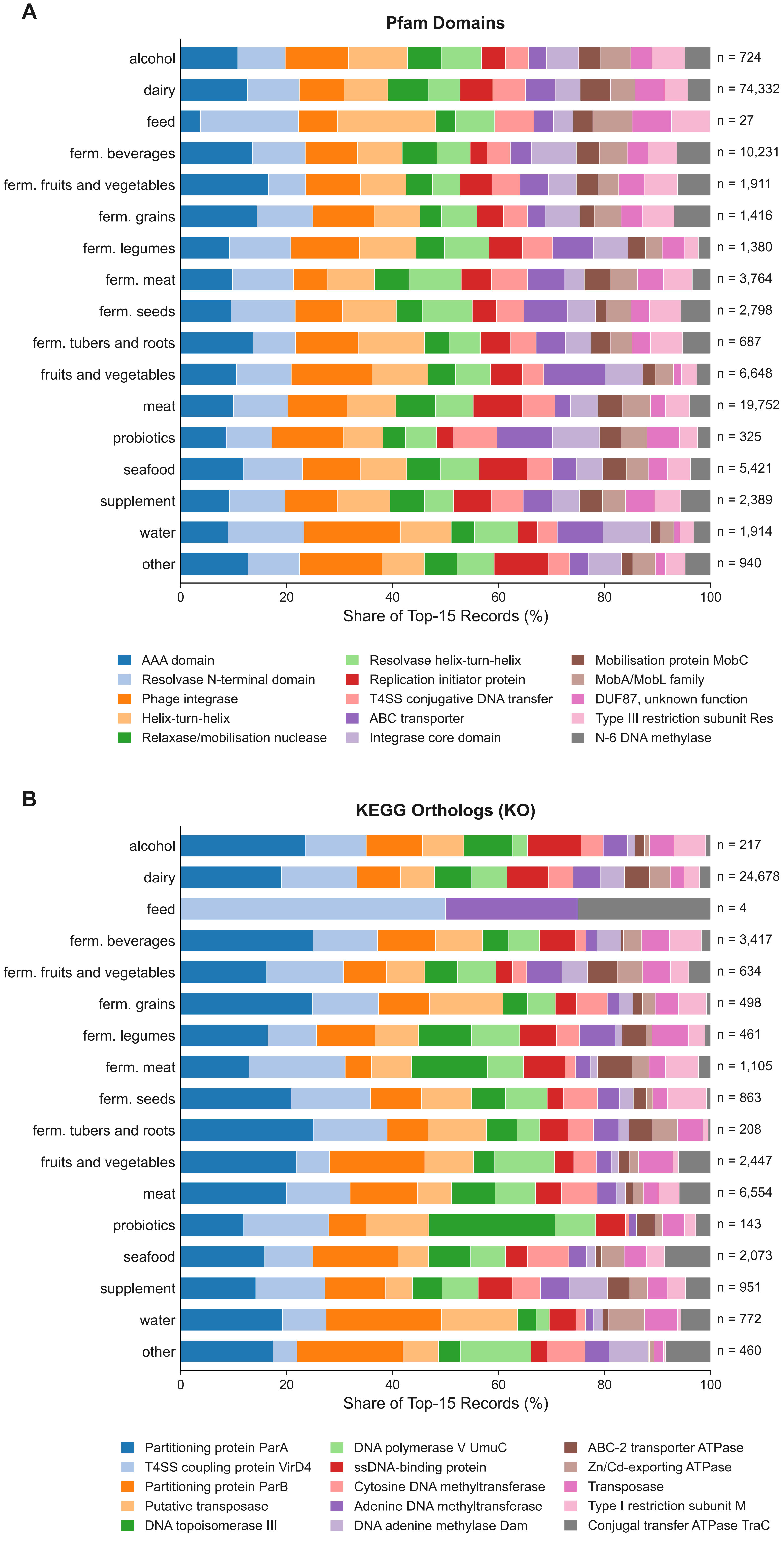
A
Pfam Domains
alcohol
dairy
feed
ferm. beverages
ferm. fruits and vegetables
ferm. grains
ferm. legumes
ferm. meat
ferm. seeds
ferm. tubers and roots
fruits and vegetables
meat
probiotics
seafood
supplement
water
other
n = 724
n = 74,332
n = 27
n = 10,231
n = 1,911
n = 1,416
n = 1,380
n = 3,764
n = 2,798
n = 687
n = 6,648
n = 19,752
n = 325
n = 5,421
n = 2,389
n = 1,914
n = 940
0
20
40
60
80
100
Share of Top-15 Records (%)
AAA domain
Resolvase N-terminal domain
Phage integrase
Helix-turn-helix
Relaxase/mobilisation nuclease
Resolvase helix-turn-helix
Replication initiator protein
T4SS conjugative DNA transfer
ABC transporter
Integrase core domain
Mobilisation protein MobC
MobA/MobL family
DUF87, unknown function
Type III restriction subunit Res
N-6 DNA methylase
B
KEGG Orthologs (KO)
alcohol
dairy
feed
ferm. beverages
ferm. fruits and vegetables
ferm. grains
ferm. legumes
ferm. meat
ferm. seeds
ferm. tubers and roots
fruits and vegetables
meat
probiotics
seafood
supplement
water
other
n = 217
n = 24,678
n = 4
n = 3,417
n = 634
n = 498
n = 461
n = 1,105
n = 863
n = 208
n = 2,447
n = 6,554
n = 143
n = 2,073
n = 951
n = 772
n = 460
0
20
40
60
80
100
Share of Top-15 Records (%)
Partitioning protein ParA
T4SS coupling protein VirD4
Partitioning protein ParB
Putative transposase
DNA topoisomerase III
DNA polymerase V UmuC
ssDNA-binding protein
Cytosine DNA methyltransferase
Adenine DNA methyltransferase
DNA adenine methylase Dam
ABC-2 transporter ATPase
Zn/Cd-exporting ATPase
Transposase
Type I restriction subunit M
Conjugal transfer ATPase TraC
